# Photobiocatalytic Diastereo- and Enantioselective Synthesis of Non-Canonical Amino Acids Bearing Challenging Stereochemical Dyads and Triads

**DOI:** 10.64898/2026.10.07.757362

**Authors:** Zhiyu Bo, Lei Cheng, Xiangyu Zhang, Amarjit Singh, Peng Liu, Yang Yang

**Author notes:** **Corresponding AuthorAmarjit Singh** − Synthetic Molecule Process Development, Takeda Development Center Americas, Inc., Cambridge, Massachusetts 02142, USA.;, **Peng Liu** − Department of Chemistry, University of Pittsburgh, Pittsburgh, Pennsylvania 15260, United States;, **Yang Yang** − Department of Chemistry and Biochemistry, University of California Santa Barbara, Santa Barbara, California 93106, United States; Department of Bioengineering, Biomolecular Science and Engineering (BMSE) Program, and Howard Hughes Medical Institute, University of California Santa Barbara, Santa Barbara, California 93106, United States. Z.B., L.C., and X.Z. contributed equally.

## Abstract

Catalyst-controlled stereodivergent synthesis of acyclic molecules bearing multiple stereogenic centers remains a longstanding challenge in asymmetric catalysis, particularly through intermolecular radical C–C coupling. Here, we report the directed evolution of pyridoxal 5’-phosphate (PLP)-dependent biocatalysts for the stereodivergent synthesis of non-canonical amino acids bearing either skipped α,γ-stereo-chemical dyads or contiguous α,β,γ-stereochemical triads. Starting from a thermophilic tryptophan synthase β-subunit, protein engineering generated complementary enzyme variants that selectively construct three of the four possible α,γ-stereoisomers through biocatalyst-controlled reversal of both remote γ-stereochemistry and proximal α-stereochemistry. These engineered enzymes catalyzed intermolecular radical C–C bond formation with a broad range of secondary alkyltrifluoroborates, affording α,γ-disubstituted amino acids in excellent yields and stereoselectivities. Furthermore, directed evolution enabled efficient β-dehydroxylative radical coupling of the non-native threonine-derived amino-acrylate intermediate, providing amino acids bearing contiguous α,β,γ-stereochemical triads with up to 96:4:<0.1:<0.1 d.r. and >99:1 e.r.. To further demonstrate the synthetic utility of this platform, the stereochemically enriched amino acids were transformed into valuable γ-chiral amines through engineered PLP-dependent tryptophan decarboxylases, establishing a dual PLP enzyme-based biocatalytic cascade for the enantiodivergent synthesis of privileged amine scaffolds. Molecular dynamics simulations reveal that enzyme-controlled γ-stereoselectivity originates from active-site remodeling that modulates π–π interactions and steric contacts governing the facial selectivity of the radical addition step. This work expands the synthetic capabilities of radical biocatalysis by enabling catalyst-controlled stereodivergent assembly of complex acyclic stereochemical arrays and establishes engineered PLP enzymes as versatile catalysts for the synthesis of stereochemically complex amino acids and chiral amines.

## INTRODUCTION

The past decade has witnessed significant progress in the stereoselective construction of contiguous stereogenic centers.^1^ Despite these notable advances, catalytic asymmetric synthesis of more complex stereo-chemical arrays, including contiguous 1,2,3-stereochemical triads and skipped 1,3-stereochemical dyads, remains underdeveloped (Scheme 1(A)). Even more challenging is the catalyst-controlled, stereodivergent synthesis of such stereochemically complex scaffolds, in which multiple stereoisomers are selectively accessed from common substrates through catalyst-regulated reversal of stereochemical outcomes.^2^ In particular, this challenge is further magnified in radical-mediated intermolecular C–C coupling, where highly reactive open-shell intermediates must be transformed within a well-defined chiral environment to suppress non-stereoselective pathways.^3,4^ In particular, compared to stereocontrol in conformationally constrained cyclic systems, the catalytic diastereo- and enantioselective synthesis of stereochemical dyads and triads in acyclic systems via intermolecular C–C coupling continues to present significant challenges.^5^

**Scheme1.**
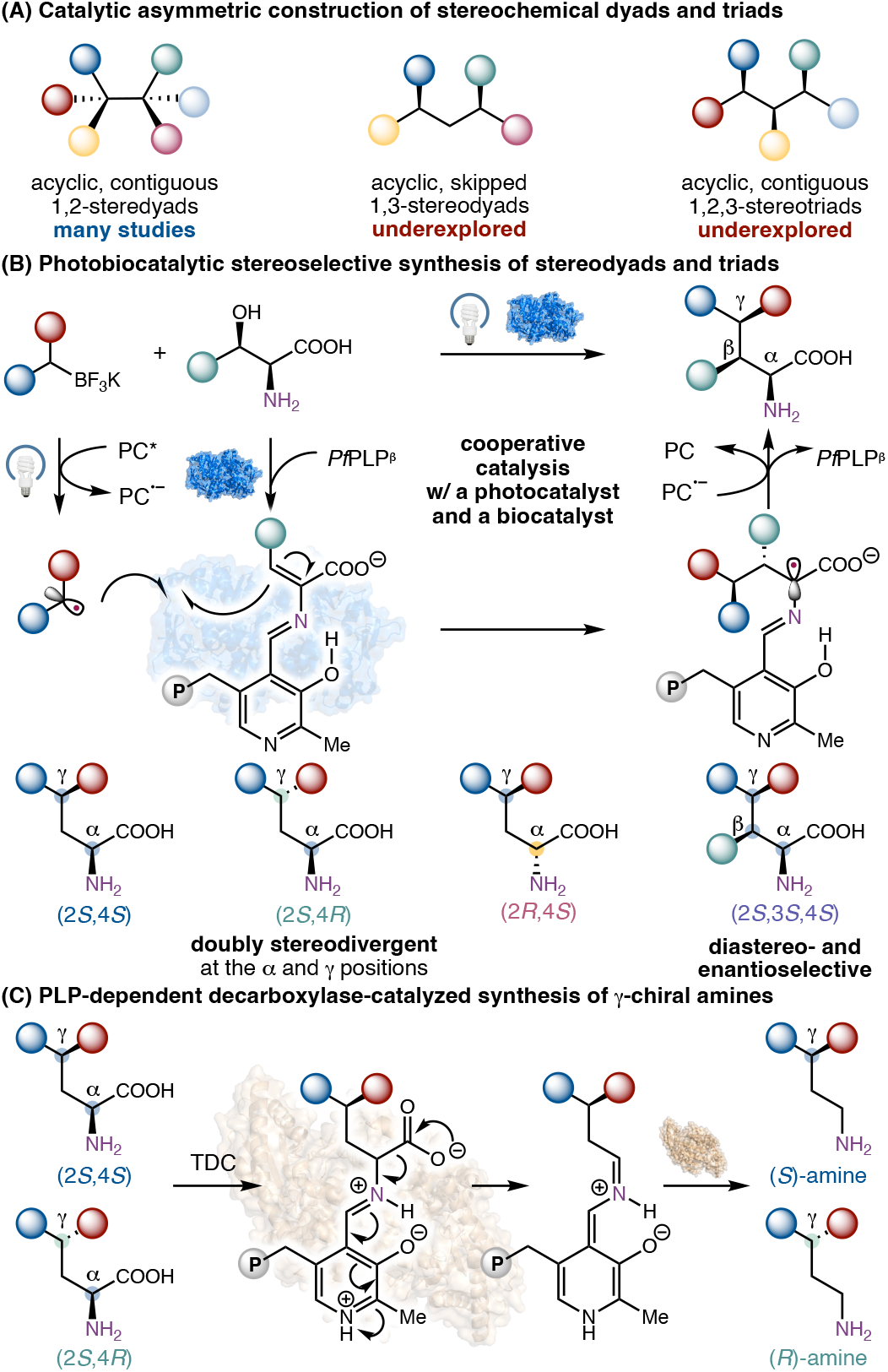
Diastereo- and Enantioselective Photobiocatalytic Synthesis of Skipped 1,3-Stereochemical Dyads and Contiguous 1,2,3-Stereochemical Triads: Overview of This Study

Enabled by the intimate substrate-protein interactions within the active site, enzymes are capable of imposing excellent stereocontrol over challenging intermolecular bond forming events, thereby providing a promising solution to otherwise difficult problems in asymmetric catalysis.^6,7^ Over the past decade, by leveraging highly reactive open-shell intermediates for C–C bond formation, new-to-nature asymmetric radical biocatalysis has emerged as a powerful strategy for the stereoselective synthesis of molecular complexity, complementing classical biocatalytic processes relying on a closed-shell mechanism.^8,9^ However, despite the groundbreaking developments in this field and the numerous stereoselective biocatalysts engineered by directed evolution, new-to-nature radical-mediated biocatalytic assembly of adjacent acyclic 1,2,3-stereogenic centers and skipped 1,3-stereogenic centers remains rare^13e,13f^ further highlighting the challenges facing biocatalysis researchers and protein engineers.

In this context, our laboratory has recently focused our efforts on the biocatalytic stereoselective synthesis of stereochemically complex non-canonical amino acids. Due to their essential roles in the development of peptide therapeutics and bioactive agents,^10^ the past several years have witnessed a renaissance of the search for efficient catalytic methods to access non-canonical amino acids in a stereochemically defined fashion.^11^ Recently, by reprogramming native two-electron pyridoxal 5’-phosphate-dependent enzymes^12^ to catalyze single-electron transformations, our group and others advanced pyridoxal radical biocatalysis as a powerful platform for the asymmetric synthesis of useful non-canonical amino acids.^13,14^ Based on our prior work repurposing tryptophan synthases^15^ for stereoselective radical C–C bond formation,^13a,13d,13g^ we envisioned that the further optimization of these PLP biocatalysts could enable the construction of a wider range of challenging amino acids bearing two to three stereocenters, including those possessing a 1,2,3-stereochemical triad and a 1,3-skipped stereochemical dyad which remained difficult to access via established methods.

To realize this proposal, several notable challenges must be addressed. First, as the γ-position is well removed from the PLP cofactor, controlling the stereochemistry at this remote site via intermolecular radical addition to the enzymatic aminoacrylate intermediate represents a non-trivial task for accessing both α,γ-stereodyads and α,β,γ-stereotriads (Schemes 1(B)). Indeed, remote stereocenter construction has long been regarded as a challenging task in organocatalysis.^16^ In particular, the doubly stereodivergent biocatalytic synthesis of α,γ-chiral amino acids at both the α- and γ-positions has remained elusive.^11, 13, 14^ Furthermore, the diastereo- and enantioselective synthesis of non-canonical amino acids featuring an α,β,γ-stereotriad would require the use of threonine, a non-native amino acid donor of tryptophan synthases. The substantially faster undesired deaminative conversion of threonine into pyruvate would further complicate efficient biocatalytic C–C bond formation (Scheme 1(B)).^17^ In this work, we present the successful development of new PLP-dependent C–C coupling biocatalysts allowing the stereocontrolled synthesis of non-canonical amino acids bearing either an α,γ-stereodyad or an α,β,γ-stereotriad. Beyond their value as non-canonical amino acids, these stereochemically complex products can serve as versatile precursors to other privileged chiral scaffolds. In particular, enzymatic decarboxylation of α,γ-substituted amino acids would provide a direct strategy for translating stereochemically enriched amino acid products into synthetically useful γ-chiral amines.^18^ To demonstrate this potential, the α,γ-chiral amino acid products generated by the photobiocatalytic platform were further subjected to biocatalytic decarboxylation, enabling enantiodivergent access to valuable enantioenriched γ-chiral amines.

## RESULTS AND DISCUSSION

At the commencement of this study, we first investigated the photobiocatalytic construction of amino acid stereochemical dyads using secondary alkyltrifluoroborate salt **1a** and serine **2a** as the model substrates using the previously engineered thermophilic tryptophan synthase β-subunit variant *Pf*PLP^β^.^17a^ It was found that *Pf*PLP^β^ exhibited low levels of activity and diastereoselectivity, giving rise to a mixture of two diastereomers (2*S*,4*S*)-**3a** and (2*R*,4*S*)-**3a** in 9% combined yield and 44:56 diastereomeric ratio (Scheme 2(A), entry 1). Despite the low initial activity, the introduction of the Y301H mutation to *Pf*PLP^β^ dramatically improved its catalytic activity and diastereoselectivity, giving rise to (2*S*,4*S*)-**3a** as the major product with 90:10 d.r. and >99:1 e.r. (entry 2). Careful stereochemical assignment through Marfey’s analysis (See the SI for details) revealed that *Pf*PLP^β^ Y301H imposed excellent stereocontrol over the remote γ-stereocenter (4*S*: 4*R* >99:1), while the stereochemical purity at the α-position of **3a** was 90:10.

**Scheme2.**
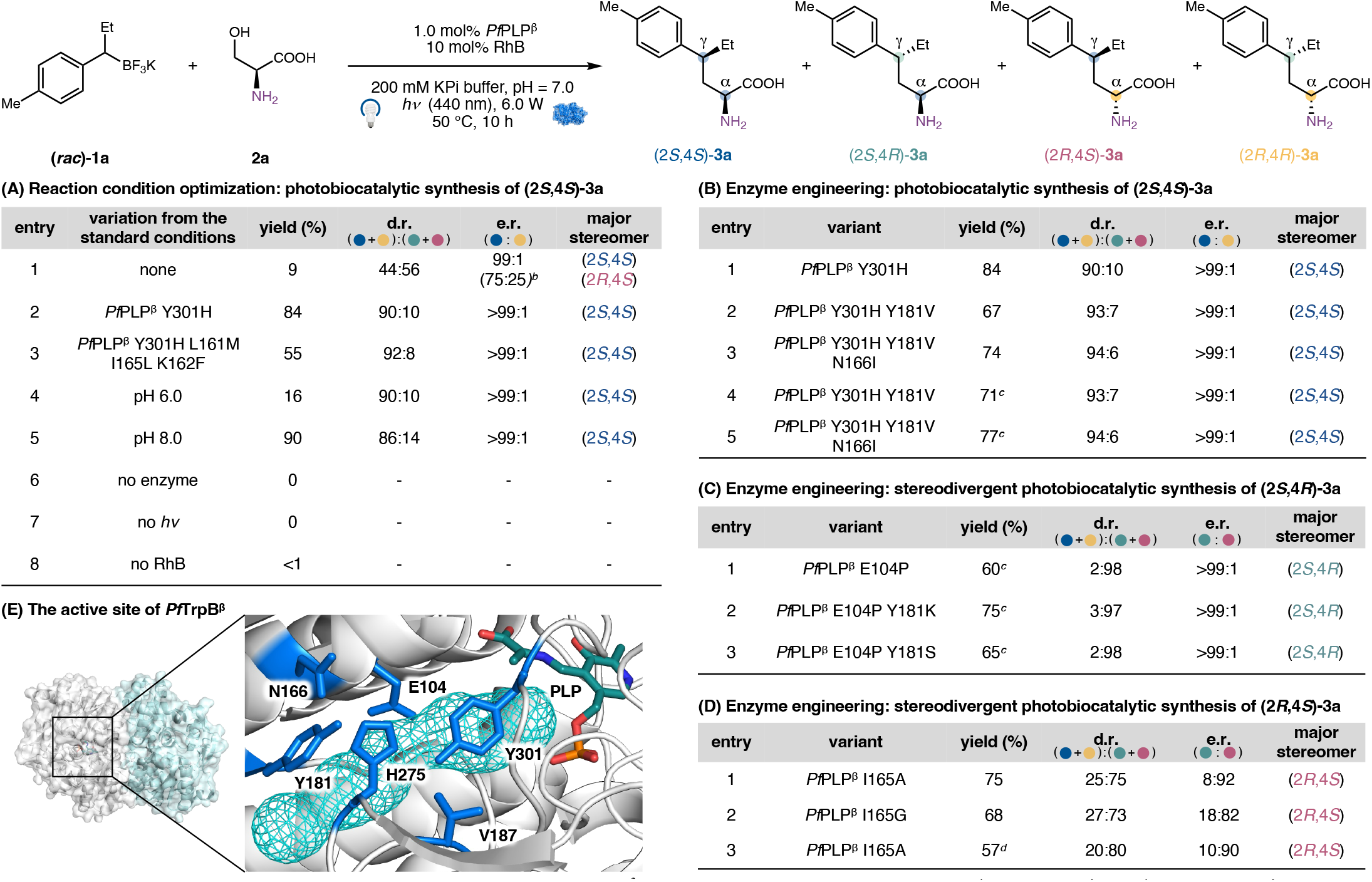
Enzyme Engineering for Stereodivergent Photobiocatalytic Synthesis of Amino Acids Bearing α,γ-Stereochemical Dyads. ^*a*^Reaction conditions: **1a** (3.3 mM), **2a** (16.4 mM), 1.0 mol% *Pf*PLP^β^ variant (33.0 μmol), 10 mol% RhB (330.0 μmol), *hν* (440 nm, 6 W), 200 mM KPi buffer (pH = 7.0), DMSO (8% v/v), 50 °C, 10 h. RhB: rhodamine B. Yields were determined by LC–MS; d.r. and e.r. were determined by product derivatization with Marfey’s reagent and UPLC–MS. The d.r. values were further confirmed by ^1^H NMR analysis. See the SI for details. ^*b*^The second major stereomer was (2*R*,4*S*); the e.r. in parentheses was (2*R*,4*S*)-**3a**:(2*S*,4*R*)-**3a**. ^*c*^1.5 mol% *Pf*PLP^β^ variant was used. ^*d*^pH 10.0 was used. (E) The active site and substrate tunnel of *Pf*PLP^β^. The substrate tunnel was shown in PyMOL with the crystal structure of *Pf*PLP^β^ (PDB ID: 5VM5).

To further improve the stereocontrol over the α position of amino acid products, we next evaluated our previously engineered PLP biocatalyst variants that showed enhanced α-stereoselectivity during the synthesis of amino acids with a single α-stereocenter (entry 3).^13d^ Although the *Pf*PLP^β 2.0^ mutant (*Pf*PLP^β^ Y301H L161M I165L K162F) showed improved diastereoselectivity, its activity was also reduced. Additionally, our reaction condition optimization studies showed that a lower pH of the reaction medium led to a higher (*S*):(*R*) ratio at the α-position of the amino acid product but decreased the overall yield (entry 4). Increasing the pH favored (*R*)-configuration at the α-position of amino acid products (entry 5). Therefore, pH 7.0 was chosen for further reaction development. Control experiments in the absence of the enzyme (entry 6) and visible light irradiation (entry 7) showed no product formation, confirming the dual photobiocatalytic nature of this transformation. Performing this reaction in the absence of rhodamine B afforded <1% yield (entry 8), indicating the essential role of this organic photocatalyst.

To enhance the diastereocontrol of this photobiocatalytic radical transformation, we carried out directed evolution using *Pf*PLP^β^ Y301H as the parent enzyme (Schemes 2(B), entry 1). Guided by substrate tunnel analysis with CAVER, ^19^ residues at the substrate tunnel or within the active site were targeted for site-saturation mutagenesis (SSM) and screening (Scheme 2(E)). Through high throughput photobiocatalysis using an in-house assembled 96-position LED array and a 96-position parallel reactor, we evaluated SSM libraries for the stereoselective synthesis of α,γ-chiral amino acids. During the first round of engineering, Y181V was identified as a beneficial mutation, increasing the diastereomeric ratio to 93:7 with a slightly reduced yield (Schemes 2(B), entry 2). Subsequent engineering revealed N166I as an additional advantageous mutation, resulting in concurrent improvements in yield and diastereoselectivity (entry 3). The use of 1.5 mol% of this triple mutant *Pf*PLP^β^ Y301H Y181V N166I (*Pf* PLP^β*S,S*-dyad^) afforded (2*S*,4*S*)-**3a** in 77% yield and 94:6 d.r., establishing *Pf* PLP^β*S,S*-dyad^ as an effective biocatalyst for the diastereoselective formation of (2*S*,4*S*)-skipped stereochemical dyads (entry 5). Next, we turned our attention to accessing the complementary stereoisomeric products through the reversal of the remote γ-stereoselectivity. Evaluation of our *Pf*PLP^β^ mutant library revealed *Pf*PLP^β^ E104P as an effective variant to invert the γ-configuration while maintaining excellent stereocontrol over the α-position with the same (2*S*)-stereopreference, furnishing (2*S*,4*R*)-**3a** in 60% with 2:98 d.r. and >99:1 e.r. (Scheme 1(C), entry 1). Interestingly, in our prior study, the E104P mutation was found to reverse the α-stereochemistry to favor the (2*R*)-configuration during the synthesis of amino acids bearing a single α-stereocenter,^13d^ suggesting a departure in stereoinduction mechanism during the construction of α,γ-stereodyad. An additional round of protein engineering via SSM and screening identified *Pf*PLP^β^ E104P Y181K as the most efficient γ-reversal variant, providing (2*S*,4*R*)-**3a** in 75% yield with 3:97 d.r. (entry 2). This variant was designated as *Pf* PLP^β*S,R*-dyad^ for subsequent substrate scope studies. Furthermore, we found that *Pf*PLP^β^ I165A and *Pf*PLP^β^ I165G led to inversion of α-stereochemistry, allowing (2*R*,4*S*)-**3a** to form in 75% yield (25:75 d.r. and 8:92 e.r.) and 68% yield (27:73 d.r. and 18:82 e.r.), respectively (Scheme 2(D), entries 1–2). Using *Pf* PLP^β^ I165A, increasing the reaction pH to 10.0 improved the stereoselectivity to 20:80 d.r. and 10:90 e.r. with a yield of 57% (entry 3). Accordingly, *Pf* PLP^β^ I165A was designated as *Pf* PLP^β*R,S*-dyad^. Together, through the engineering of the common parent PLP enzyme *Pf*PLP^β^, we achieved the stereodivergent synthesis of non-canonical amino acids bearing α,γ-stereodyads by inverting both the α,γ-stereocenters.

With the optimal stereodivergent variants established, we next examined the substrate scope for the photobiocatalytic synthesis of non-canonical amino acids bearing 1,3-skipped stereocenters (Table 1). It was found that *Pf* PLP^β^_*S,S*-dyad_ showed excellent diastereo- and enantiocontrol over the synthesis of the (2*S*,4*S*)-amino acids with diverse substitution patterns. Under the optimized reaction conditions, the parent substrate provided **3a** in 77% yield with excellent stereocontrol (96:4 d.r., >99:1 e.r.). Electron-donating substituents such as a *para*-methoxy (**1c**) group and electron-neutral substituents such as a *para*-methylthio (**1d**) group were readily accommodated, affording the corresponding products with excellent diastereo- and enantioselectivity (93:7 d.r. and >99:1 e.r.). In addition, halogenated organotrifluoroborate substrates bearing a fluoro (**1e**), a chloro (**1f**), and a bromo (**1g**) substituent were also efficiently transformed, delivering the corresponding amino acid products in good yield and excellent stereoselectivity. The presence of chlorine and bromine functional group handles would allow further derivatization to be carried out using established cross-coupling technologies.^20^ Organoboron substrates possessing a *meta*-substituted was also compatible, providing (2*S*,4*S*)-**3h** in 73% yield with 94:6 d.r. and >99:1 e.r. Finally, a pyridyl substrate was also transformed into (2*S*,4*S*)-**3i** in 47% yield with 90:10 d.r. and >99:1 e.r., demonstrating the compatibility of this photobiocatalytic method with stereoselective syntheses of medicinally relevant *N*-heterocycles. ^21^ The absolute configuration of **3b** was determined by derivatization of the product obtained with *Pf* PLP^β^_*S,S*-dyad_, followed by comparison of its chiral HPLC chromatogram with that reported in the literature (see the SI for details).

**Table 1.** Substrate Scope of Photobiocatalytic Stereodivergent Synthesis of α, γ, -Amino Acid Stereochemical Dyads. ^*a*^Reaction conditions for *Pf* PLP^β^_*S,S*-dyad_: (*rac*)-1 (3.3 mM), **2a** (16.4 mM), 1.5 mol% *Pf* PLP^β^_*S,S*-dyad_ (50.0 μM), 10 mol% RhB (330.0 μM), *hν* (440 nm, 6.0 W), 200 mM KPi buffer (pH = 7.0), DMSO (11% v/v), 50 °C, 10 h. Yields were determined by LC–MS; d.r. and e.r. values were determined through product derivatization with Marfey’s reagent and UPLC–MS analysis. The d.r. values were further confirmed by ^1^H NMR analysis. See the SI for details. Reaction conditions for *Pf* PLP^β^_*S,R*-dyad_ : (*rac*)-1 (3.3 mM), **2a** (16.4 mM), 1.5 mol% *Pf* PLP^β^_*S,R*-dyad_ (50.0 μM), 10 mol% RhB (330.0 μM), *hν* (440 nm, 2.0 W), 200 mM KPi buffer (pH = 7.0), DMSO (11% v/v), 50 °C, 10 h. ^*b*^DMSO (16% v/v) was used. ^*c*^1.5 mol% *Pf* PLP E104P was used.

| (2*S*,4*S*)-**3** | (*rac*)-**1** | **2a** | (2*S*,4*R*)-**3** |  |  |
| --- | --- | --- | --- | --- | --- |
| (2*S*,4*S*)-**3a**<br>(76 ± 2)% yield<br>95:5 d.r.<br>>99:1 e.r. | (2*S*,4*S*)-**3b**<br>(77 ± 1)% yield<br>94:6 d.r.<br>>99:1 e.r. | (2*S*,4*S*)-**3c**<br>(78 ± 1)% yield<br>93:7 d.r.<br>>99:1 e.r. | (2*S*,4*R*)-**3a**<br>(75 ± 2)% yield<br>3:97 d.r.<br>>99:1 e.r. | (2*S*,4*R*)-**3b**<br>(86 ± 1)% yield<sup>c</sup><br>21:79 d.r.<br>82:18 e.r. | (2*S*,4*R*)-**3c**<br>(88 ± 1)% yield<br>[106 mg scale, 77% yield]<br>2:98 d.r., >99:1 e.r. |
| (2*S*,4*S*)-**3d**<br>(36 ± 2)%<sup>b</sup> yield<br>93:7 d.r.<br>>99:1 e.r. | (2*S*,4*S*)-**3e**<br>(80 ± 1)%<sup>b</sup> yield<br>94:6 d.r.<br>>99:1 e.r. | (2*S*,4*S*)-**3f**<br>(74 ± 2)%<sup>b</sup> yield<br>92:8 d.r.<br>>99:1 e.r. | (2*S*,4*R*)-**3d**<br>(62 ± 1)%<sup>b</sup> yield<br>4:96 d.r.<br>>99:1 e.r. | (2*S*,4*R*)-**3e**<br>(80 ± 1)% yield<sup>b</sup><br>5:95 d.r.<br>99:1 e.r. | (2*S*,4*R*)-**3f**<br>(80 ± 1)% yield<sup>b</sup><br>3:97 d.r.<br>>99:1 e.r. |
| (2*S*,4*S*)-**3g**<br>(47 ± 1)%<sup>b</sup> yield<br>91:9 d.r.<br>>99:1 e.r. | (2*S*,4*S*)-**3h**<br>(73 ± 1)% yield<br>94:6 d.r.<br>>99:1 e.r. | (2*S*,4*S*)-**3i**<br>(47 ± 2)% yield<br>90:10 d.r.<br>>99:1 e.r. | (2*S*,4*R*)-**3g**<br>(71 ± 1)% yield<sup>b</sup><br>3:97 d.r.<br>>99:1 e.r. | (2*S*,4*R*)-**3h**<br>(70 ± 1)% yield<br>21:79 d.r.<br>86:14 e.r. | (2*S*,4*R*)-**3i**<br>(75 ± 2)% yield<br>6:94 d.r.<br>>99:1 e.r. |

The substrate scope of the stereocomplementary (2*S*,4*R*)-variant *Pf* PLP^β^_*S,R*-dyad_ closely mirrored that of *Pf*PLP^β^_*S,S*-dyad_. It was found that organotrifluoroborate substrates bearing electron-donating (**1c**), electron-neutral (**1d**) and electron-withdrawing (**1e–1g**) groups were well tolerated, affording products in good yield with excellent diastereo- and enantioselectivities. *Pf*PLP^β^_*S,S*-dyad_ exhibited reduced diastereo- and enantioselectivity for unsubstituted (**1b**) and *meta*-substituted (**1h**) substrates but retained excellent activity. The relative and absolute stereochemistry of **3c** prepared by *Pf* PLP^β^_*S,R*-dyad_, was further confirmed by single-crystal X-ray diffraction analysis of **3c**’’ prepared from **3c** via *N*-Boc protection and amidation (Scheme 3; see the SI for details). Moreover, the photobiocatalytic transformation could be readily scaled up to 100 mg-scale synthesis without loss of diastereo and enantiopurity using commercially available Kessil LED lamps (440 nm), thereby further showcasing the synthetic utility of our methods.

**Scheme3.**
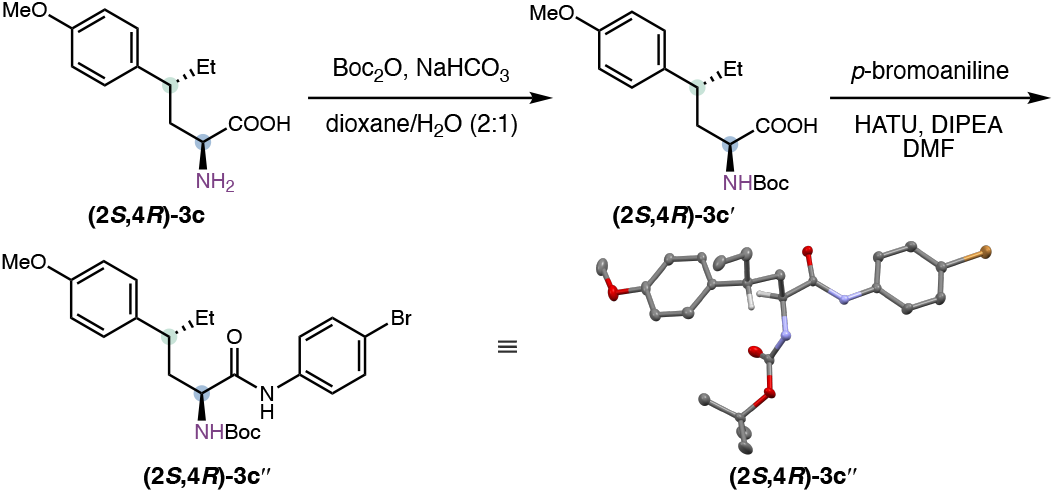
Relative and Absolute Stereochemistry Determination of 1,3-Stereodyads Produced by *Pf* PLP^β^_*S,R*-dyad_. ^*a*^See the SI for detailed reaction conditions. For x-ray crystal structures, thermal ellipsoids were set at 50% probability; hydrogen atoms are omitted for clarity.

Together, the diastereo- and enantiocomplementary enzyme variants *Pf* PLP^β^_*S,S*-dyad_, *Pf* PLP^β^_*S,R*-dyad_ and *Pf* PLP^β^_*R,S*-dyad_ constitute a set of engineered biocatalysts for the doubly stereodivergent synthesis of amino acids bearing skipped α,γ-stereocenters. To further exemplify this capability of our methods, we transformed **1c** into three stereoisomers (Scheme 4). Under our optimized reaction conditions, *Pf*PLP^β^_*S,S*-dyad_ furnished (2*S*,4*S*)-**3c** in 78% yield with 93:7 d.r. and >99:1 e.r.. *Pf* PLP^β^_*S,R*-dyad_ reversed the γ-stereochemistry to provide (2*S*,4*R*)-**3c** in 88% yield with 2:98 d.r. and >99:1 e.r.. Finally, *Pf* PLP^β^_*R,S*-dyad_ inverted the α-stereocenter, affording (2*R*,4*S*)-**3c** in 87% yield with 13:87 d.r. and 5:95 e.r..

**Scheme4.**
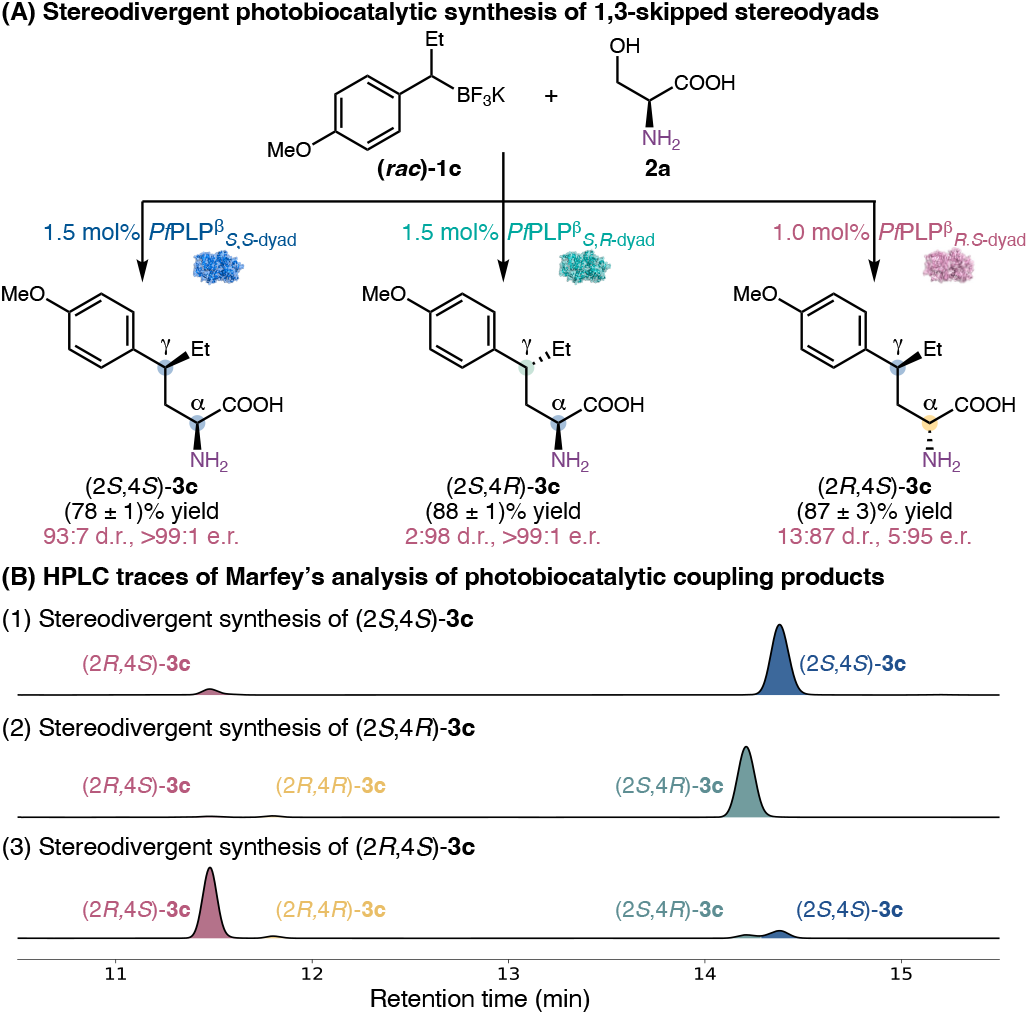
Diastereo- and Enantioselective Photobiocatalytic Synthesis of Skipped 1,3-Stereochemical Dyads

Next, we turned our attention to the stereoselective photobiocatalytic synthesis of non-canonical amino acids bearing contiguous α,β,γ-stereocenters, thereby further increasing the stereochemical complexity of the amino acid products. As threonine is not the native amino acid substrate of tryptophan synthases, the efficient β-dehydroxylative radical coupling of threonine is more challenging than that of serine. Our initial studies on the photobiocatalytic coupling of threonine with 1-arylethyltrifluoroborate substrates with an aromatic substituent showed diminished yields and diastereoselectivities when *Pf*PLP^β^ was applied. For example, when organoboron substrate **4a** was used, *Pf*PLP^β^ provided **5a** in only 53% yield with a moderate diastereomeric ratio of 87:8:4:1, although the major diastereomer still formed with excellent enantioselectivity (>99:1 e.r.). To improve the activity and diastereoselectivity of radical coupling enzymes, we initiated a protein engineering campaign using both acyclic and cyclic secondary alkyltrifluoroborate salts **4a** and **4i** as the substrates. For the biotransformation of **4a** (Schemes 5(A)), our first round of SSM and screening targeting substrate tunnel residues identified Y301F as a beneficial mutation, improving the yield of **5a** to 65% and the diastereomeric ratio to 90:7:3:<0.1. Further engineering revealed H275E as an additional advantageous mutation, leading to a further increase in enzyme activity and diastereoselectivity (72% yield, 92:3:4:1 d.r. and > 99:1 e.r.). The resulting double mutant, *Pf*PLP^β^ Y301F H275E, was designated as *Pf*PLP^β^_triad 1_.

**Scheme5.**
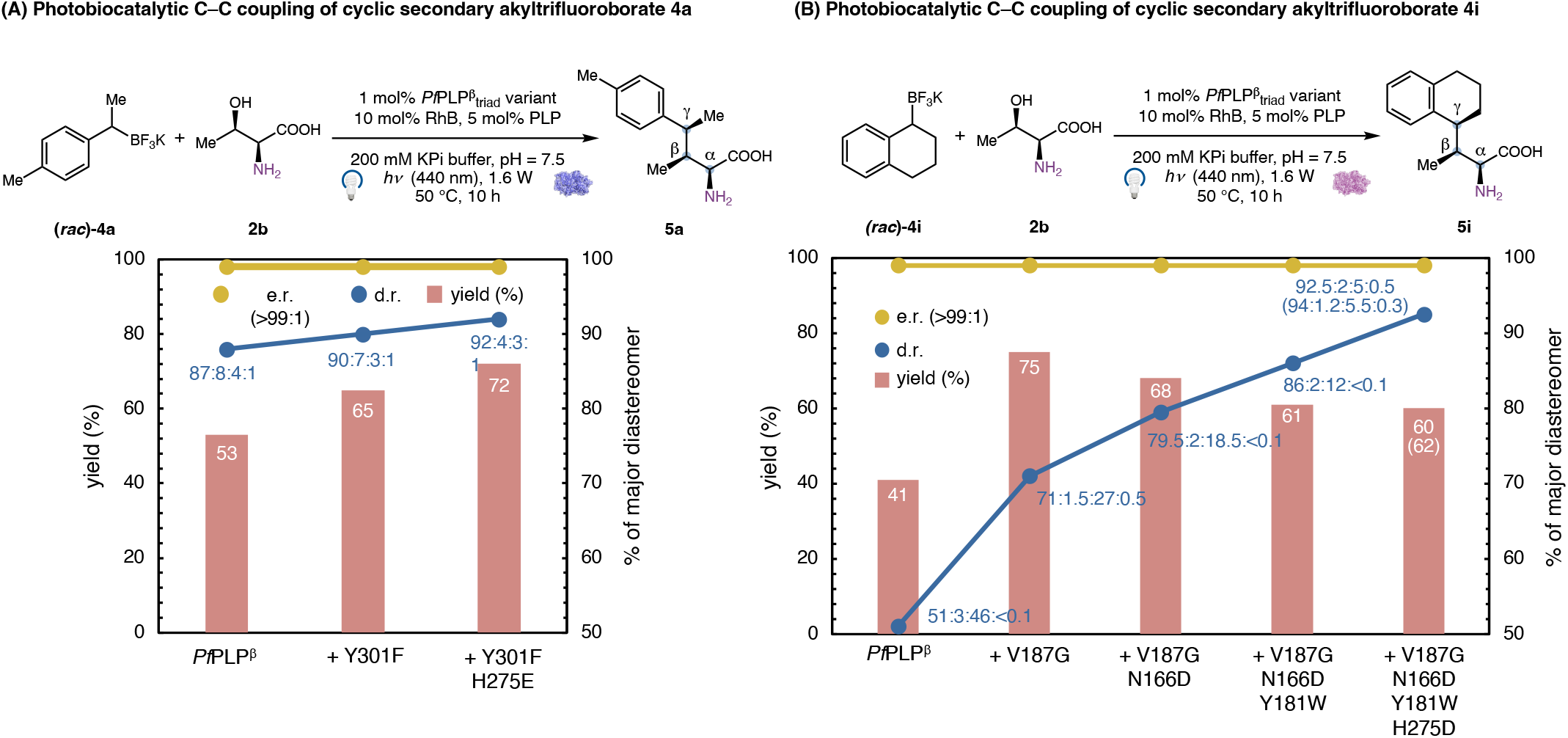
Directed Evolution for the Diastereo- and Enantioselective Photobiocatalytic Synthesis of α,β,γ-Stereochemical Triads. ^*a*^Reaction conditions: (*rac*)-**4a** or (*rac*)-**4i** (5.6 mM), **2b** (28.0 mM), 1.0 mol% *Pf*PLP^β^_triad_ variant, 10 mol% RhB, 5 mol% PLP, *hν* (440 nm, 1.6 W), 200vmM KPi buffer (pH = 7.5), DMSO (19% v/v), 50 °C, 10 h. Yields were determined by LC–MS; d.r. and e.r. were determined through product derivatization with Marfey’s reagent and UPLC–MS analysis. The d.r. values were further confirmed by ^1^H NMR analysis. See the SI for details.

When cyclic organotrifluoroborate substrate **4i** was used, a further reduced yield and diastereoselectivity were observed with *Pf*PLP^β^ _triad 1_ (41% yield, 51:3:46:<0.1 d.r., >99:1 e.r., Schemes 5(B)). Furthermore, the use of *Pf*PLP^β^ _triad 1_ and its intermediate variant did not afford improved activity and diastereoselectivity. To develop effective biocatalysts for the transformation of these cyclic substrates, we re-evolved *Pf*PLP^β^ using **4i** as the substrate. The first round of engineering identified V187G as a pivotal mutation, affording **5i** in 75% yield, 71:1.5:27:0.5 d.r. and >99:1 e.r.. Subsequent rounds of protein engineering aimed at enhancing the diastereoselectivity of the enzyme revealed N166D and Y181W as beneficial mutations, allowing **5i** to form with 86:2:12:<0.1 d.r.. Finally, the incorporation of H275D led to a quadruple mutant, *Pf*PLP^β^ V187G N166D Y181W H275D (*Pf*PLP^β^ _triad 2_), affording **5i** in 61% yield with 92.5:2:5:0.5 d.r. and >99:1 e.r.. Increasing the enzyme loading to 1.5 mol% led to slight improvement in yield and diastereomeric ratio (62% yield, 94:1.2:5.5:0.3 d.r. and >99:1 e.r.). The absolute stereochemistry of **5k** prepared by *Pf* PLP^β^ V187G N166D Y181M, was further confirmed by single-crystal X-ray diffraction analysis (see the SI for details). These results established *Pf*PLP^β^_triad 2_ as an effective biocatalyst for the stereoselective construction of amino acid bearing adjacent α,β,γ-stereocenters from cyclic organotrifluoroborate substrates. Collectively, the *Pf*PLP^β^ _triad 1_ and *Pf*PLP^β^_triad 2_ lineages provided a panel of useful enzyme variants for the diastereo- and enantioselective photobiocatalytic synthesis of amino acid stereotriads.

Next, we examined the substrate scope for the photobiocatalytic synthesis of amino acids with three contiguous stereocenters (Table 2). Un-substituted substrate **4b** provided desired product **5b** in 85% yield with an excellent 95:3:2:<0.1 d.r. and >99:1 e.r.. Additionally, a range of *para*-substituents on the aromatic ring, including a fluoro (**4c**), a chloro (**4d**), a bromo (**4e**), or a methoxy (**4f**) group, were also tolerated. The sterically demanding *ortho*-substituted substrate **4h** was also successfully converted, delivering the desired product with 92:4:4:<0.1 d.r. and >99:1 e.r., although with a slightly reduced yield. In addition to acyclic secondary alkyltrifluoroborate substrates, cyclic organoboron substrates including a tetrahydronaphthalene (**5i**) and a chromane (**5j**) were also efficiently transformed with excellent diastereo- and enantiocontrol. Five-membered secondary alkylboron reagents such as an indanyl (**4k**) substrate were also readily accommodated. Furthermore, a bulky benzofuran substrate was also successfully converted to the desired amino acid product (5l) in 64% yield with 93:5.5:1:0.5 d.r. and >99:1 e.r..

**Table 2.**
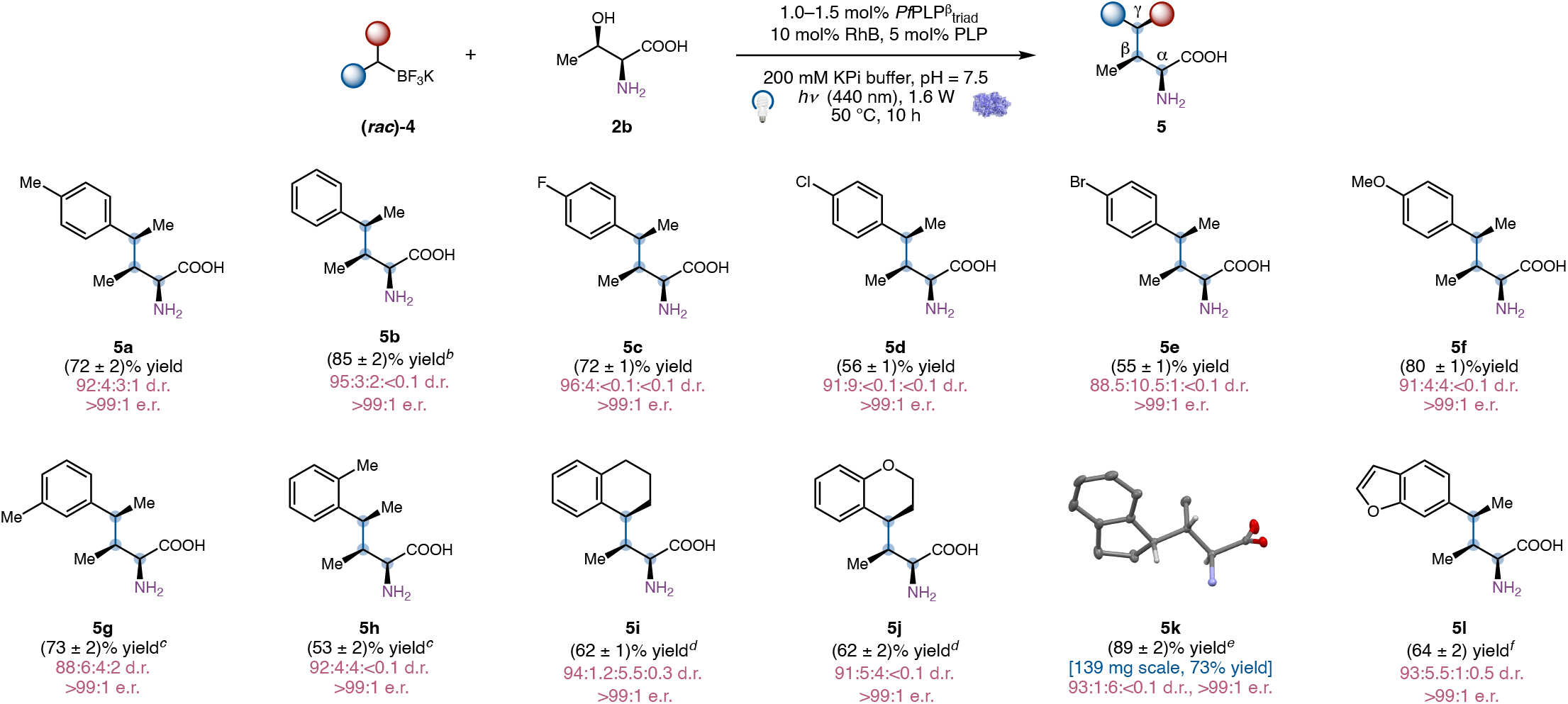
Substrate Scope of the Photobiocatalytic Stereoselective Synthesis of α,β,γ-Amino Acid Stereochemical Triads. ^*a*^Reaction conditions: (*rac*)-**4** (5.6 mM), **2b** (28.0 mM), 1.0 mol% *Pf* PLP^β^_triad 1_(56.0 μmol), 10 mol% RhB (560.0 μmol), 5 mol% PLP (280.0 μmol), *hν* (440 nm, 1.6 W), 200 μM KPi buffer (pH = 7.5), DMSO (19% v/v), 50 °C, 10 h. Yields were determined by LC–MS analysis; d.r. and e.r. values were determined through product derivatization with Marfey’s reagent and UPLC–MS analysis. The d.r. values were further confirmed by ^1^H NMR analysis. See the SI for details. ^*b*^1.0 mol% *Pf* PLP^β^, DMSO (14% v/v) was used. ^*c*^1.0 mol% *Pf* PLP^β^ V187G N166D was used. ^*d*^1.5 mol% *Pf* PLP^β^_triad 2_ was used. ^*e*^1.0 mol% *Pf* PLP^β^ V187G N166D Y181M was used. ^*f*^1.5 mol% *Pf* PLP^β^ V187G N166D Y181W was used.

With the ability to conveniently access non-canonical amino acids bearing multiple stereocenters, we next questioned whether we could transform these amino acid products into the corresponding γ-chiral amines using engineered PLP-dependent decarboxylases (Scheme 6). However, due to the bulky nature of non-canonical amino acids derived from our photobiocatalytic C–C coupling methods, their biocatalytic decarboxylation represented a nontrivial task. Toward this objective, we evaluated several PLP-dependent tyrosine decarboxylases^22^ and tryptophan decarboxylases^23^ for the conversion of **3e** to **6**. While tyrosine decarboxylase from *Enterococcus faecium* (*Efi*1TyrDC)^22b^ did not affect this challenging amino acid decarboxylation (Schemes 6(A), entry 1), *Ruminococcus gnavus* tryptophan decarboxylase exhibited promising activity, likely due to the similar geometry of non-canonical amino acid **3** with tryptophan. In particular, a previously engineered triple mutant *Rgn*TDC H120N L126M W349Y (*Rgn*TDC^NMY^)^23b^ showed decarboxylation activity for both (2*S*,4*S*)-**3e** and (2*S*,4*R*)-**3e**, with the (2*S*,4*R*)-diastereomer displaying higher activity (entry 2). Further optimization of reaction parameters including the reaction buffer, pH and temperature improved the yield of this biocatalytic decarboxylation to 85% (entry 3). To further enhance the catalytic efficiency of *Rgn*TDC^NMY^ toward larger non-canonical amino acid substrates, guided by the AutoDock model of the PLP external aldimine derived from (2*S*,4*R*)-**3e**, we carried out site-directed mutagenesis and assayed the activity of these *Rgn*TDC^NMY^ mutants. These studies led to the identification of T356S as a beneficial mutation, allowing the preparation of (*R*)-**6** in 89% yield and >99:1 e.r. (entry 3). The T356S mutation also facilitated the decarboxylation of the mismatched diastereomer (2*S*,4*S*)-**3e**, delivering the opposite enantiomer (*S*)-**6** in 85% yield and >99:1 e.r. (*S*:*R*) with 2.5 mol% biocatalyst loading (entry 3). Together, the successful development of this photobiocatalytic C–C coupling/tryptophan decarboxylase-catalyzed amino acid decarboxylation cascade afforded a new strategy for the asymmetric synthesis of γ-chiral amines from easily available organoboron reagents, thereby further demonstrating the excellent synthetic potential of biocatalytic cascades using dual PLP-dependent enzymes (Schemes 6(B)).

**Scheme6.**
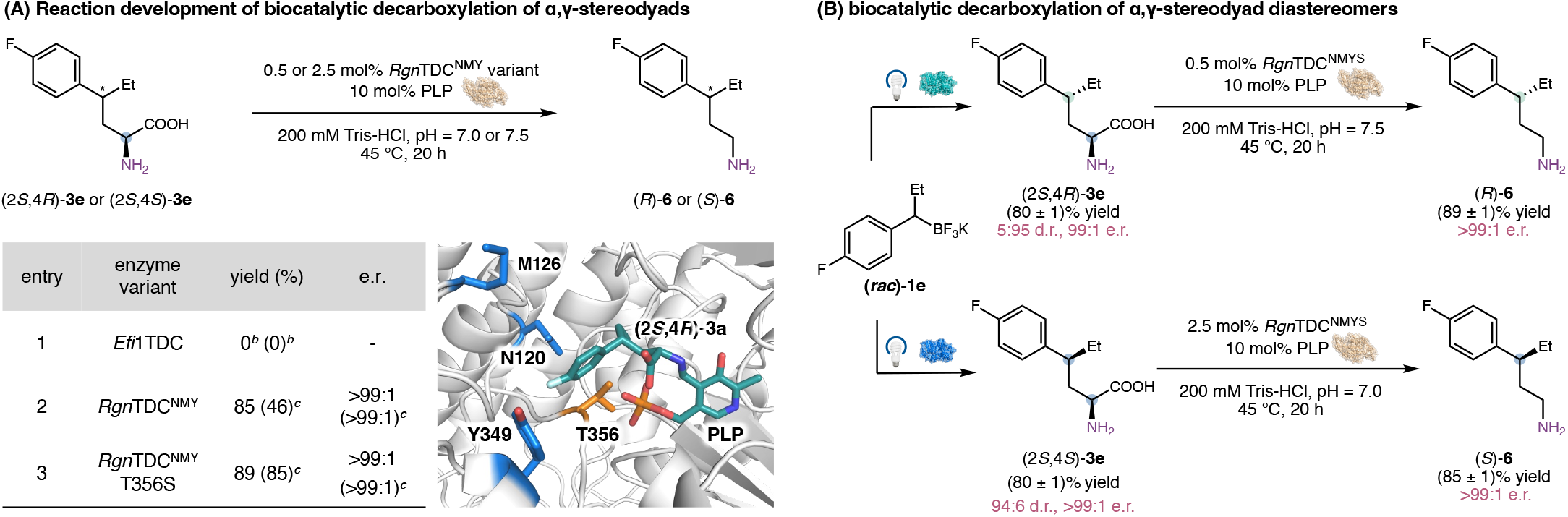
Biocatalytic Enantiodivergent Decarboxylation of α,γ-Chiral Amino Acids to γ-Chiral Amines. ^*a*^Reaction conditions: (2*S*,4*R*)-**3e** (7.5 mM), 0.5 mol% *Rgn*TDC^NMY^ variant (37.5 μM), 10 mol% PLP (750.0 μmol), 200 mM Tris-HCl buffer (pH = 7.5), 45 °C, 20 h. Yields were determined by LC–MS; the e.r. (*R*:*S*) of (*R*)-**6** was determined through product derivatization and chiral HPLC. ^*b*^Reaction conditions: (2*S*,4*R*)-**3e** or (2*S*,4*S*)-**3e** (7.5 mM), 0.5 mol% *Efi*1TDC (37.5 μM), 10 mol% PLP (750.0 μmol), 200 mM KPi buffer (pH = 5.5), 25 °C, 20 h. ^*c*^Results in parentheses are yields and e.r. (*S*:*R*) of (*S*)-**6** derived from (2*S*,4*S*)-**3e** as the substrate. Reaction conditions: (2*S*,4*S*)-**3e** (7.5 mM), 2.5 mol% *Rgn*TDC^NMY^ variant (187.5 μM), 10 mol% PLP (750.0 μmol), 200 mM Tris-HCl buffer (pH = 7.0), 45 °C, 20 h. Docking model was generated via AutoDock with the crystal structure of *Rgn*TDC^NMY^ (PDB ID: 4OBV).

To gain insights into the origin of γ-stereoselectivity in the C−C bond-forming radical addition step, we performed classical molecular dynamics (MD) simulations to investigate key interactions between active-site residues and the incoming secondary alkyl radical (Scheme 7, see the SI for computational details). Since the tryptophan synthase β-subunit variant *Pf*PLP^β^ functions as a homodimer in solution,^24^ a dimeric model was used in our MD simulations. To elucidate the origin of the enzyme-controlled stereodivergence in *Pf*PLP^β^_*S,S*-dyad_- and *Pf*PLP^β^_*S,R*-dyad_-catalyzed C−C coupling of serine (**2a)** and secondary alkyl trifluoroborate 1c, we modeled the near-attack conformations (NACs) of the γ-stereochemistry-determining radical addition of 1-(4-methoxyphenyl)propyl radical (**1c′)** derived from **1c** to the C_*β*_ of the serine-derived aminoacrylate intermediate. Three independent 1000 ns replicas were performed to model the two competing pathways exposing either the (*Re*)- or (*Si*)-face of the secondary alkyl radical **1c′** to the enzymatic aminoacrylate by restraining the 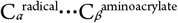 distance to 2.3−2.6 Å using a harmonic potential of 200 kcal·mol^−1^·Å^−2^ and the forming bond angle and dihedral angles to better resemble the radical addition transition state geometries (see Figure S1 for details).

**Scheme7.**
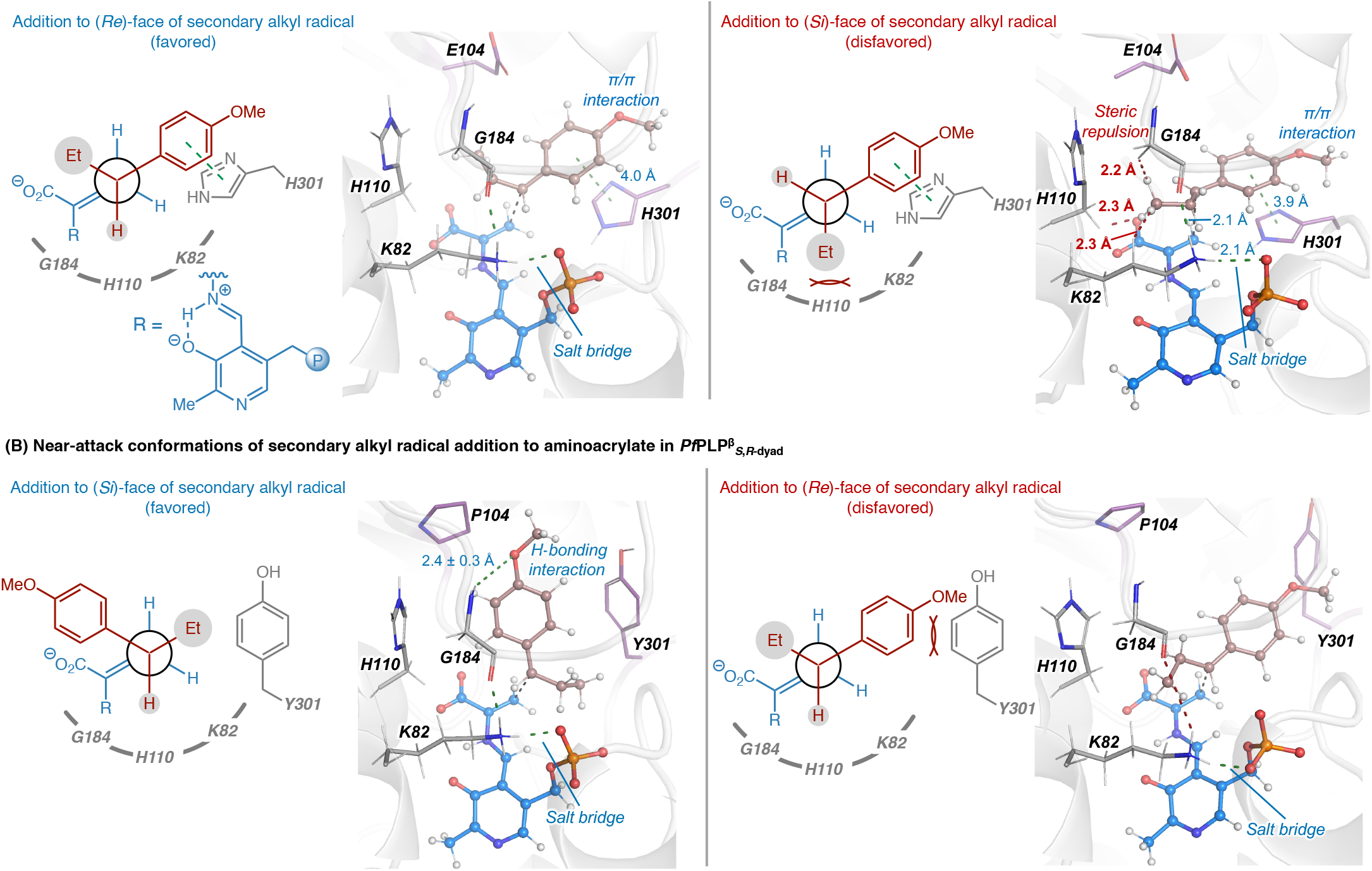
Computational Studies on Photobiocatalytic Radical Coupling: Understanding the Origin of Enzymatic γ-Stereocontrol in the C–C Bond-Forming Radical Addition Step Using Classical Molecular Dynamics (MD) Simulations. ^*a*^Distances in the figures are median values during the NAC simulations.

In the NAC simulations with the *Pf*PLP^β^_*S,S*-dyad_ variant (Schemes 7(A)), a persistent π/π interaction between the side chain of H301 and the 4-methoxyphenyl group of the secondary alkyl radical was observed in both (*Re*)- and (*Si*)-radical addition pathways (π^radical^···π^H301^ = 4.0 and 3.9 Å, respectively, see Figure S10 for details). This π···π interaction places the 4-methoxyphenyl group *anti*-periplanar to the aminoacrylate C_*α*_*=*C_*β*_ double bond (see the Newman projections in Schemes 7(A)). In the (*Re*)-face addition pathway, this orientation places the smallest H atom on the benzylic carbon towards the salt bridge between K82 side chain and the PLP phosphate. By contrast, in the (*Si*)-face addition, the larger Et group points towards K82, leading to steric clashes with K82 and surrounding residues including H110 and G184. Besides the K82– phosphate salt bridge, a persistent hydrogen bonding interaction between G184 and K82 (O^G184^···H _Namine_^K82^ = 2.1 Å) further restricts the motion of these residues and accentuates the steric clashes with the Et group. These results suggest that the stereoselectivity in the radical addition with *Pf*PLP^β^_*S,S*-dyad_ is controlled by both π/π interaction with H301 and steric contacts with K82 and surrounding residues that disfavor the (*Si*)-face addition pathway.

The NAC simulations with the *Pf*PLP^β^_*S,R*-dyad_ variant revealed substantially different active-site residue interactions with the incoming secondary alkyl radical from those with the *Pf*PLP^β^_*S,S*-dyad_ variant (Schemes 7(B)). The π···π interaction with H301 is disrupted when this site is replaced with a bulkier Y301 residue, which does not engage in stabilizing π···π interaction with the 4-methoxyphenyl group of the alkyl radical. As a result, in the (*Si*)-face addition pathway, the 4-methoxyphenyl group is placed in a *gauche*^*−*^ position with respect to the aminoacrylate C_*α*_*=*C_*β*_ double bond. This orientation avoids the steric clash with the bulky Y301 residue and is further stabilized by a hydrogen bonding interaction between the 4-methoxyl group and the main chain of G184. In addition, the E104P mutation creates more space in the active site to better accommodate the bulky 4-methoxyphenyl group to occupy the *gauche*^*−*^ position proximal to the smaller P104 residue. In the (*Re*)-face addition NACs with *Pf*PLP^β^_*S,R*-dyad_, the secondary alkyl radical adopts the same *anti* conformation observed with the *Pf*PLP^β^_*S,S*-dyad_ variant. However, the lack of stabilizing π···π interaction combined with the steric clash with the bulky Y301 residue leads to larger fluctuations of Y301 and contributes to the divergent stereoselectivity that disfavors this (*Re*)-face addition pathway. Collectively, these results indicate *Pf*PLP^β^_*S,R*-dyad_’s preference for (*Si*)-face radical addition is due to the substantially modified active site steric environment that disrupts the π···π interaction with residue 301 in the *anti*-conformation and creates more space to accommodate the bulky 4-methoxyphenyl group in the *gauche*^*−*^ conformation in the (*Si*)-face addition pathway.

## CONCLUSION

In summary, this study established a stereodivergent photobiocatalytic platform for the synthesis of stereochemically complex non-canonical amino acids through radical-mediated intermolecular C–C bond formation. Through the directed evolution of tryptophan synthase variants, engineered PLP-dependent biocatalysts were developed for the highly diastereo- and enantioselective construction of both skipped α,γ-stereochemical dyads and contiguous α,β,γ-stereochemical triads. In particular, biocatalyst-controlled stereochemical reversal enabled the doubly stereodivergent synthesis of amino acids bearing α,γ-stereocenters, allowing selective access to three of the four possible stereoisomers from common substrates. Our photobiocatalytic methods further accommodated the synthesis of amino acids containing three contiguous stereocenters with excellent stereocontrol across a broad substrate scope. Mechanistic and computational studies provided insights into how engineered active-site environments govern radical addition stereoselectivity and remote stereocontrol. Beyond amino acid synthesis, the resulting α,γ-chiral amino acids were further transformed into valuable enantioenriched γ-chiral amines through newly engineered PLP-dependent decarboxylases, thereby expanding the synthetic utility of these photobiocatalytic coupling products. Collectively, this study highlighted the power of radical pyridoxal biocatalysis for controlling challenging contiguous and remote stereochemical elements in open-shell intermolecular bond-forming reactions, providing a versatile strategy for selectively accessing stereochemically complex building blocks that remain difficult to prepare through conventional catalytic methods.

## ASSOCIATED CONTENT

### Supporting Information

Experimental procedures, computational details, DNA and protein sequences, characterization data, HPLC traces and NMR spectra (PDF).

The Supporting Information is available free of charge on the ACS Publications website.

## AUTHOR INFORMATION

### Author

**Zhiyu Bo** − Department of Chemistry and Biochemistry, University of California Santa Barbara, Santa Barbara, California 93106, United States

**Lei Cheng** − Department of Chemistry and Biochemistry, University of California Santa Barbara, Santa Barbara, California 93106, United States

**Xiangyu Zhang** − Department of Chemistry, University of Pittsburgh, Pittsburgh, Pennsylvania 15260, United States

## ACKNOWLEDGMENT

We are grateful to the National Institutes of Health (R01EB036084 to Y.Y. for biocatalysis), National Science Foundation (CHE-2400087 to P.L. for computational study) and Takeda Pharmaceuticals (A.S. and Y.Y.) and for financial support. Y.Y. is an Alfred P. Sloan Research Fellow (FG2024-22244), a Camille Dreyfus Teacher-Scholar Awardee (TC-25-084), a David & Lucile Packard Fellow (2023-76169), and an HHMI Freeman Hrabowski Scholar. Computations were carried out at the University of Pittsburgh Center for Research Computing and Data and the Advanced Cyberinfrastructure Coordination Ecosystem: Services & Support (ACCESS) Program, supported by NSF award numbers OAC-2117681, OAC-1928147, OAC-1928224 and CHE-260005.

## Table of Contents

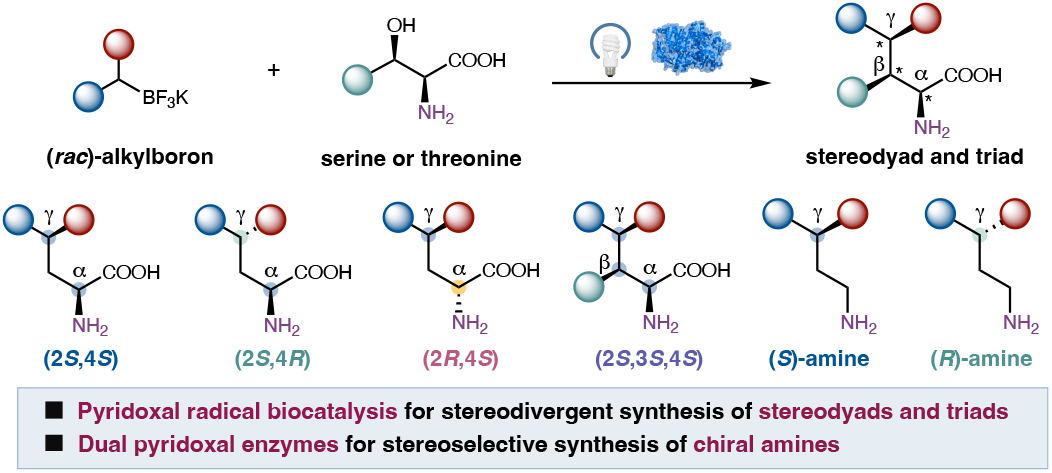

